# Deficiency in reverse cholesterol transport in mice augments sepsis

**DOI:** 10.1101/2020.04.26.051250

**Authors:** Qian Wang, Ling Guo, Dan Hao, Misa Ito, Kai-jiang Yu, Rui-tao Wang, Chieko Mineo, Philip W. Shaul, Xiang-An Li

**Affiliations:** Department of Physiology and Saha Cardiovascular Research Center, University of Kentucky, Lexington, KY 40536; Department of Pediatrics, University of Texas Southwestern Medical Center, Dallas, TX 75390; Department of Intensive Care Unit, Harbin Medical University Cancer Hospital, Harbin Medical University, Harbin, Heilongjiang 150081, China; Department of Intensive Care Unit, The First Affiliated Hospital of Harbin Medical University, Harbin Medical University, Harbin, Heilongjiang 150081, China; Department of Internal Medicine, Harbin Medical University Cancer Hospital, Harbin Medical University, Harbin, Heilongjiang, 150081, China

**Keywords:** SR-BI, Scarb1, sepsis, reverse cholesterol transport, hemoglobin

## Abstract

**Background:** Sepsis claims over 215,000 lives and costs $16.7 billion per year in America alone. Recent studies revealed that HDL receptor scavenger receptor BI (SR-BI or Scarb1) plays a critical protective role in sepsis. Using Scarb1I179N mice, a mutant SR-BI mouse model with 90% depletion in hepatic SR-BI, we previously reported that the mutant mice are susceptible to cecal ligation and puncture (CLP)-induced sepsis. However, using a hypo-AlbCreSR-BIfl/fl mouse model, Huby’s group showed that the liver-specific SR-BI KO mice are not more susceptible to CLP-induced sepsis. In this study, we generated a new floxed SR-BI mouse model to clarify the contribution of hepatic SR-BI in sepsis. SR-BI is known as a receptor that plays a key role in reverse cholesterol transport (RCT) by uptaking cholesterol to the liver. So, our established AlbCreSR-BIfl/fl mice (liver-specific SR-BI KO) is an RCT deficiency mice model that can be used to understand the mechanisms of RCT protecting against sepsis and may provide new insight into the pathogenesis of sepsis.

**Methods and Results:** We generated SR-BIfl/fl mice by flanking exon 2. We bred the floxed mice with AlbCre mice to generate AlbCreSR-BIfl/fl mice (liver-specific SR-BI KO mice), then the mice were backcrossed to C57BL/6J for 10 generations. As shown in Fig 1, the liver SR-BI expression was normal in SR-BIfl/fl mice as compared to C57BL/6J (B6) mice, but completely depleted in AlbCreSR-BIfl/fl mice. Using this liver-specific SR-BI KO model, we observed that a deficiency in RCT rendered the mice highly susceptible to CLP-induced sepsis as shown by 80% and 14.3% survival of SR-BIfl/fl and AlbCreSR-BIfl/fl mice, respectively. We found aggravated inflammatory cytokine production, altered leukocyte recruitment and slightly increased in the blood and peritoneal bacteria. Moreover, we found RCT deficiency mice increased both free and total cholesterol levels in serum and showed severer hemolysis in AlbCreSR-BIfl/fl mice than SR-BIfl/fl mice during CLP-induced sepsis. Importantly, when we fed AlbCreSR-BIfl/fl mice with probucol to decrease the cholesterol level in serum before performing CLP, the survival rate of AlbCreSR-BIfl/fl mice improved to 88.9%.

**Conclusions:** Deficiency RCT resulting in abnormal metabolism of cholesterol and lipid metabolism is a risk factor in sepsis and maintain normal metabolism of cholesterol may provide a new insight for sepsis therapies.

## Introduction

Sepsis is a life-threatening dysregulated host response associated with multiorgan hypoperfusion and dysfunction caused by infection which claims over 215,000 lives and costs $16.7 billion per year in America alone (1,2). Sepsis is still a leading cause of death in the intensive care unit and the mortality rate is about 17% to 26% in the hospital(3). The pathogenesis of sepsis is complex which is characterized by an early pro-inflammatory phase or “cytokine storm” usually causing early death. Patients who survive early sepsis often subsequent enter an immunosuppressive status and develop secondary infections(4). The current treatment mainly comprises of supportive care, rapidity administration of antimicrobials, fluid resuscitation, vasoactive medications, mechanical ventilation and nutritional support, etc.(5). The molecular mechanism is still needed to be explored to find new insight for sepsis treatment.

HDL receptor Scavenger receptor BI (SR-BI or Scarb1) is a transmembrane protein highly expressed in liver and steroidogenic tissues(6,7), such as adrenal glands, testes and ovaries(8). It is also expressed in macrophages(9), vascular smooth muscle cells and endothelial cells(10). SR-BI involving in the uptake of cholesterol esters from high-density lipoprotein (HDL) plays a key role in reverse cholesterol transport (RCT) by removing blood and excessive tissue cholesterol to the liver, protecting blood vessels and preventing the formation of atherosclerosis(11–13). Studies have revealed a crucial role of SRBI in protecting sepsis both using lipopolysaccharides (LPS) induced sepsis model(14,15) and using cecal ligation and puncture (CLP) induced sepsis model (16). SRBI whole-body knockout mice show much more susceptible to septic death. Macrophage SRBI via TLR4 signaling modulate immune response during sepsis(16) and adrenal SRBI deficiency mice fail to generate inducible corticosteroid (iCS) leading to aberrant inflammatory response during sepsis(17). We have reported that hepatic SR-BI significantly protects against CLP induced sepsis using Scarb1^I179N^ mice which is a mutant SR-BI mouse model with 90% depletion in hepatic SR-BI. Mice with 90% depletion of SR-BI in the liver cause a 3.5-fold increase in CLP septic fatality(18). However, using a hypo-AlbCreSR-BIfl/fl mouse model, Huby’s group shows that the liver-specific SR-BI KO mice are not more susceptible to CLP-induced sepsis(19). Cholesterol is the main lipid participating in sepsis which increases by 41% when mice were injected with LPS(20) and SRBI expression in the liver plays a key role in reverse cholesterol transport to maintain cholesterol metabolic homeostasis(21). But the mechanism of SR-BI involved reverse cholesterol transport in sepsis is still unclear.

In this study, we generated a new floxed SR-BI mouse model by flanking exon 2. We bred the floxed mice with AlbCre mice to generate AlbCreSR-BIfl/fl mice (reverse cholesterol transport deficiency mice) whose SRBI expression in liver is completely deleted. Using this mouse model to clarify the controversy of hepatic SR-BI in sepsis on one hand, and to explore RCT impact on sepsis on the other. Our findings revealed reverse cholesterol transport is a critical protective factor in sepsis. RCT deficiency increases serum free cholesterol levels and makes red blood cell membrane stiffen(22) inducing more serious hemolysis during sepsis. Thus, alleviation of hemolysis may be a novel mechanism of SR-BI protecting against sepsis. Probucol used to decrease serum cholesterol level shows a significant protective effect on sepsis survival.

### Experimental procedures

#### Materials and methods

Anti-SR-BI was from sigma Genosys using a 15-amino acid peptide from the C-terminal of human SR-BI. The alanine aminotransferase (ALT) kit was from BQKits, Inc. (San Diego, CA). The blood urea nitrogen (BUN) kit was from the QuantiChrom. The Cholesterol kit was from Wako. Hemoglobin assay kit was from Sigma (MAK115)

#### Animals

We generated SR-BIfl/fl mice by flanking exon 2. We bred the floxed mice to AlbCre mice to generate AlbCreSR-BIfl/fl mice (liver-specific SR-BI KO mice, the mice were backcrossed to C57BL/6J for 10 generations). Albumin-Cre mice were bought from Jackson Lab (stock NO: 003574). Mice were fed with standard laboratory rodent died. Animal experiments were approved by Animal Care and use Committee of University of Kentucky.

#### CLP induced sepsis model

CLP was performed on about 3-month-old mice as previous described(16). Mice were anesthetized and ligated at a half distance of the cecum and punched twice with 23G needle. After surgery, 1 ml phosphate-buffered saline was injected to resuscitate. For probucol administration, mice were fed with a 0.5% probucol diet (probucol was dissolved in ethanol and sprayed on normal rodent diet and dried thoroughly) for 3 days before performing CLP. The diets continued during the survival observation.

#### PCR genotyping of AlbcreSRBIfl/fl mice

Albcre genotyping primers: 5’-TGC AAA CAT CAC ATG CAC AC; 5’-TTG GCC CCT TAC CAT AAC TG and 5’-GAA GCA GAA GCT TAG GAA GAT GG. A 2% gel is required to separate bands. 351 bp for control mice, 390 bp and 351 bp for heterozygote mice.

#### Western blot

Western blot was performed as described previously(14). 30mg liver was lysed in lysis buffer with proteinase inhibitor (sigma), then after centrifugation and the supernatant was mixed with reduced buffer. Heated the mixture before loading it to the polyacrylamide gel.

#### Leukocyte recruitment to peritoneum

AlbCreSR-BI^fl/fl^ and SR-BI^fl/fl^ mice were treated with CLP for 4h and 5ml PBS were injected to the peritoneal. Count the cell number and 10^6^ cells from peritoneal fluid were used for flow cytometry. Using CD16/32 antibody (clone 93, biolegend) block against the FcR to eliminate the nonspecific staining. And then cells were incubated with anti-body Ly-6C-FITC, Ly-6G-PE from biolegend and CD11b-PerCP-5.5 and CD45-APC from BD bioscience for 30min at 4°C. The stained cells were washed with FACS buffer and analyzed by LSRII Flow Cytometer, BD. Peritoneal neutrophils (PMN, CD11b^hi^ Ly6C^hi^) and inflammatory monocytes (IM, CD11b^int^ Ly6C^hi^) were gated by CD11b and Ly6C expression on CD45^+^ cells.

#### Biochemical assays

Mice blood was collected from the abdominal aorta at 4h and 20h following CLP, and the serum is store at −80C for biochemical assays. Plasma cytokines were analyzed by Mouse Cytokine Array / Chemokine Array 31-Plex (MD31) which was provided by Eve Technologies.

#### Bacterial load

Blood and peritoneal fluid from CLP mice were diluted (20, 200 and 2000 times with dH2O), and 100ul of the dilution was plated on the LB-Agar plate and incubated at 37°C for 24h and then the number of clones was counted.

#### Statistical analysis

Survival rate was analyzed by the Kaplan-Meier method with log-rank x^2^ test. Using SPSS software to do statistical analysis of data. Data were presented as Means ± SEM. 2-tailed Student’s t-test was used to compare two groups.

Comparing more than two groups was analyzed by one-way ANOVA. Significant differences were considered at p-value <0.05.

## Results

### Reverse cholesterol transport is a protective factor in CLP-induced septic death

We confirmed the mice’s genotype by PCR. As shown in Fig.1A, SR-BIfl/fl showed one band at 351 bp and AlbCreSR-BIfl/fl mice showed multiple bands at ~390 bp and 351 bp (an artifact band of ~150 bp is frequently present from the mutant allele). The western blotting analysis showed SRBI expression in the liver of AlbCreSR-BIfl/fl mice was completely deleted. And SRBI expression in liver was similar between SRBIfl/fl mice with C57BL/6J (B6) mice (Fig.1B), proving rationality of using SR-BIfl/fl as control. As shown in Fig.1C, CLP induced 20% fatality in SR-BIfl/fl mice and 85.7% fatality in AlbCreSR-BIfl/fl mice, indicating the significant protective role of hepatic SRBI in reducing septic death. To evaluate CLP-induced sepsis effect on organ injury between the two groups of mice, kidney injury was assessed by measuring serum blood urea nitrogen (BUN) levels and liver injury was assessed by measuring serum alanine aminotransferase (ALT) levels. Kidney injury and liver injury showed little difference between AlbCreSR-BIfl/fl mice and SR-BIfl/fl mice at both 4h and 20h post CLP (Fig. D and E). Thus, mice with reverse cholesterol transport deficiency were more susceptible to CLP-induced septic death.

**Fig. 1.**
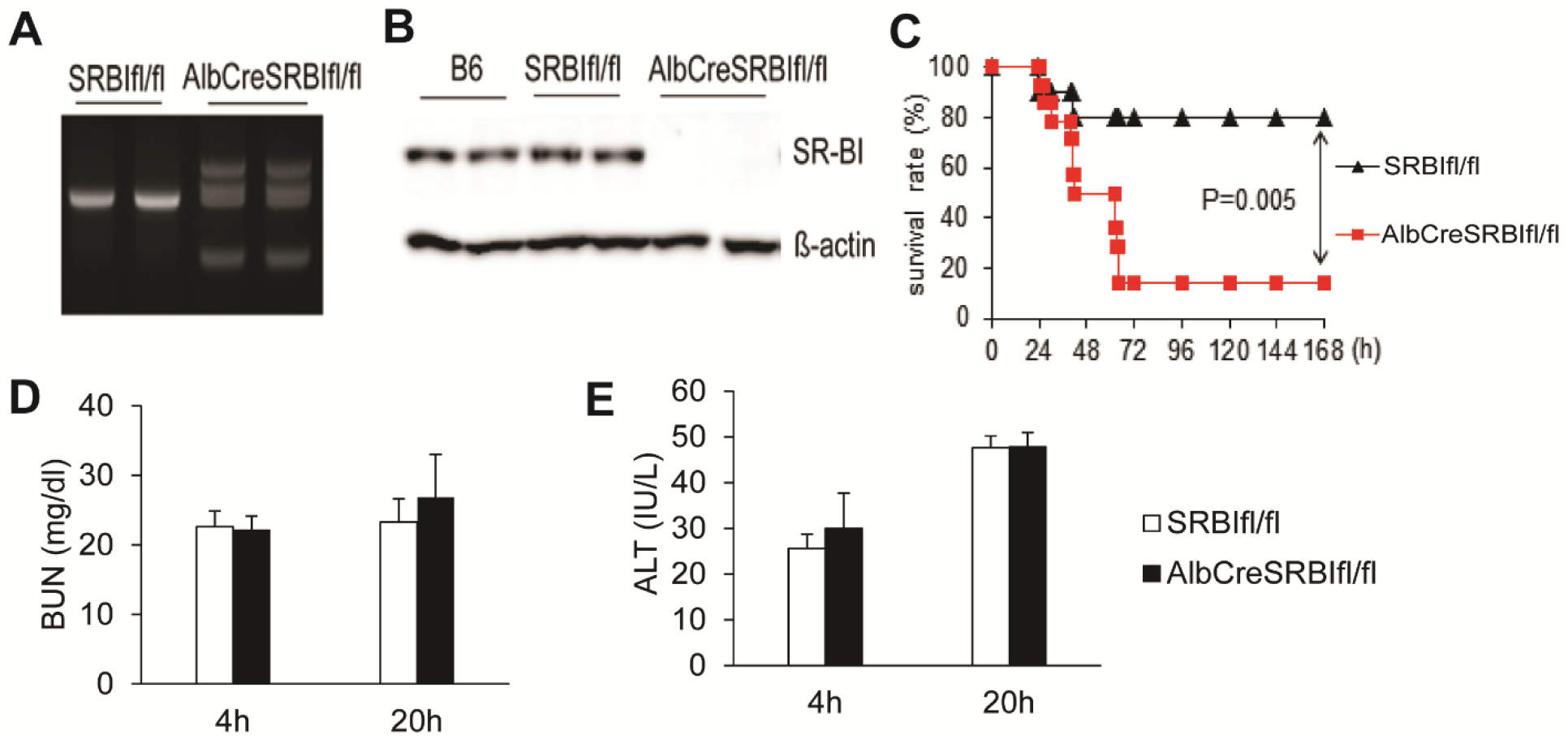
Mice deficient in hepatic SR-BI are susceptible to CLP-induced septic death. A, PCR genotyping of AlbCreSR-BI^fl/fl^ mice using tail DNA. B, Western blot analysis of SR-BI expression in the liver; C, survival analysis. AlbCreSR-BI^fl/fl^ (n = 14) and SR-BI^fl/fl^ mice (n = 10) were treated with CLP (23G needle, half ligation) and survival was observed for 7 days. The data were expressed as the percentage of mice surviving at indicated times, and survival was analyzed by Log-Rank *x*^2^ test. D, AlbCreSR-BI^fl/fl^ and SR-BI^fl/fl^ mice were treated with CLP for the indicated times, and kidney injury was assessed by measuring blood urea nitrogen (BUN) levels; and E, liver injury was assessed by measuring serum alanine aminotransferase (ALT) levels. n = 6 - 7 each group, mean ± SEM.

### Deficient RCT Exacerbates Immune Response and neutrophils recruitment during sepsis

To understand the cause of AlbCreSR-BIfl/fl mice susceptible to septic death, we assessed 31 plasma cytokine/chemokine production at 4h and 20h after CLP. AlbCreSR-BI^fl/fl^ mice had higher Eotaxin and LIX production than control mice at 20h after CLP (Fig.2A and B). IL-6 and TNF-α also showed slightly elevated in AlbCreSR-BIfl/fl mice compared to SR-BIfl/fl mice (Fig.2 C and D). Other cytokines levels showed no significant because of the big variation (Supplemental Table S1). To quantified the living bacteria in the peritoneum and in circulation, the blood and peritoneal fluid were used for analyzing bacteria load. The deficiency of hepatic SR-BI slightly increased blood bacteria and peritoneal bacteria at CLP 20h (Fig.2 E and F). Peritoneal leukocyte recruitment was analyzed by flow cytometry (Fig2.G). AlbCreSR-BI^fl/fl^ and SR-BI^fl/fl^ mice peritoneal fluid were collected at CLP 4h and then peritoneal neutrophils (PMN, CD11b^hi^ Ly6C^hi^) and inflammatory monocytes (IM, CD11b^int^ Ly6C^hi^) were gated by CD11b and Ly6C expression on CD45^+^ cells. A similar cell number of IMs cells was detected in AlbCreSR-BI^fl/fl^ and SR-BI^fl/fl^ mice but more recruited PMNs were detected in CLP-induced AlbCreSR-BI^fl/fl^ mice than SR-BI^fl/fl^ mice. Deficiency of reverse cholesterol transport induced more neutrophils recruitment to the peritoneal cavity. Above all, deficiency of RCT moderately affects bacteria in blood and peritoneal but increased leukocyte recruitment to the peritoneal cavity in sepsis.

**Fig. 2.**
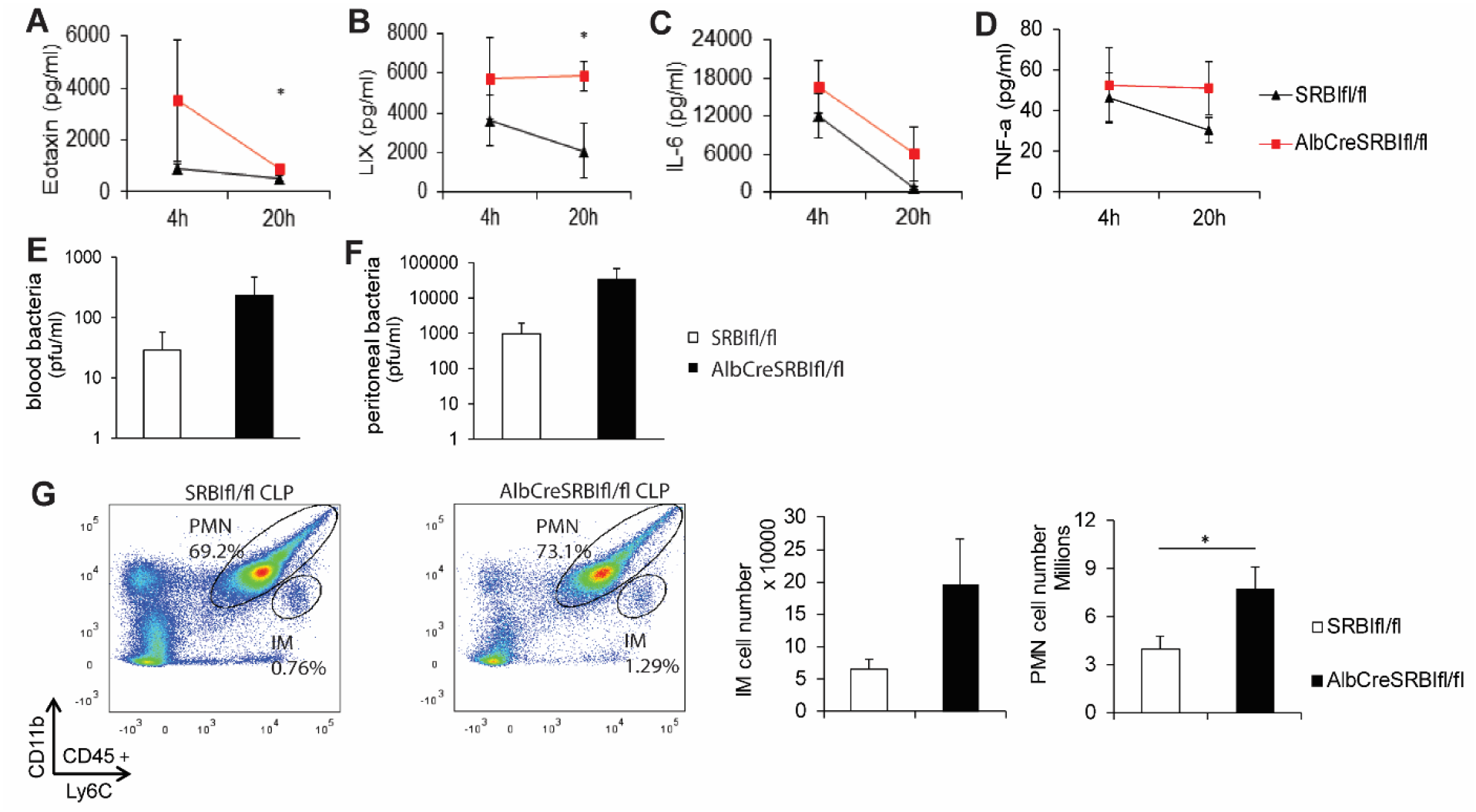
Exacerbated serum inflammatory cytokine production and more leukocyte recruitment to peritoneal cavity in AlbCreSR-BI^fl/fl^ mice during sepsis. AlbCreSR-BI^fl/fl^ and SR-BI^fl/fl^ mice were treated with CLP (23G needle, half ligation) for 4 and 20h, and the serum was analyzed for cytokines level. A and B, serum eotaxin and LIX levels. C and D, serum IL-6 and TNF-a levels. E and F, AlbCreSR-BI^fl/fl^ and SR-BI^fl/fl^ mice were treated with CLP for 20h and the blood and peritoneal fluid was analyzed for bacteria load. Deficiency of RCT slightly affects blood bacteria and peritoneal bacteria at CLP 20h. G, AlbCreSR-BI^fl/fl^ and SR-BI^fl/fl^ mice were treated with CLP for 4h and peritoneal fluid was collected. Peritoneal neutrophils (PMN, CD11b^hi^ Ly6C^hi^) and inflammatory monocytes (IM, CD11b^int^ Ly6C^hi^) were gated by CD11b and Ly6C expression on CD45^+^ cells. A similar cell number of IMs in CD45^+^ cells was detected in AlbCreSR-BI^fl/fl^ and SR-BI^fl/fl^ mice and more recruited PMNs were detected in CLP-induced AlbCreSR-BI^fl/fl^ mice than SR-BI^fl/fl^ mice. 2-tailed Student’s t-test was used to compare two groups. n = 6 - 7 each group, mean ± SEM. *p<0.05 versus SR-BI^fl/fl^ mice.

### Reverse cholesterol transport deficiency causes higher free cholesterol and total cholesterol levels and more serious hemolysis during sepsis

Neither immune response nor bacteria load in the organ is enough to explain the septic death of AlbCreSR-BI^fl/fl^ mice. The death of AlbCreSR-BI^fl/fl^ mice may have a different mechanism than our previous study found. In SR-BI^fl/fl^ mice, total cholesterol and free cholesterol levels tended to decrease at 4h (total cholesterol 68.6 ± 6.6 mg/dl; free cholesterol 12.1 ± 1.7 mg/dl) and increase at 20h (total cholesterol 116.1 ± 8.9 mg/dl; free cholesterol 37.4 ± 2.1 mg/dl) post CLP (Fig. 3 A and B). The same tendency was also observed in AlbCreSR-BI^fl/fl^ mice with higher levels of cholesterol than SR-BI^fl/fl^ mice at the designated CLP time point. But the free cholesterol level became extremely high at CLP 20h due to RCT deficiency when compared with SR-BI^fl/fl^ mice. As shown in Fig. B, free cholesterol level in AlbCreSR-BI^fl/fl^ mice was 5.5-fold higher than SR-BI^fl/fl^ mice at 20h post CLP. Cholesterol ester levels were calculated by the value difference between total cholesterol and free cholesterol (Fig. 3 C). The deficiency of RCT causes accumulated free cholesterol may affect red blood cell membrane stability and causes hemolysis during sepsis. AlbCreSR-BI^fl/fl^ showed 1.9-fold higher hemoglobin levels than SR-BI^fl/fl^ mice at 20h after CLP (Fig. 3E). Hemoglobin levels in AlbCreSR-BI^fl/fl^ during sepsis give us an insight that hemolysis may be a mechanism that causes RCT deficiency mice death during sepsis.

**Figure 3.**
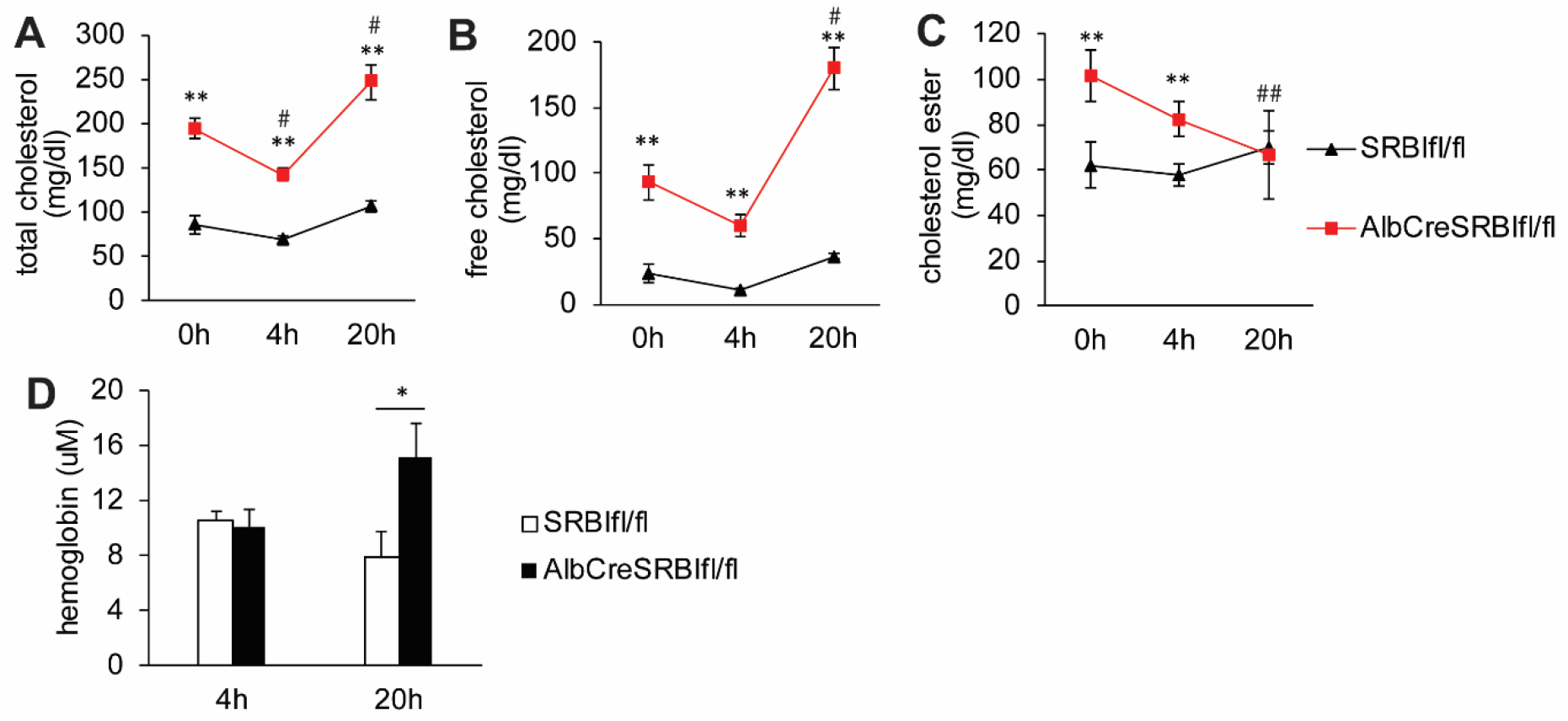
Deficiency of RCT causes hemolysis in CLP induced sepsis. AlbCreSR-BI^fl/fl^ and SR-BI^fl/fl^ mice were treated with CLP (23G needle, half ligation) A, the serum total cholesterol levels and B, the free cholesterol levels at 4h and 20h post CLP. C, the serum hemoglobin level was analyzed at 4h and 20h after CLP. 2-tailed Student’s t-test was used to compare two groups. Comparing more than two groups was analyzed by two-way ANOVA. n = 6 each group, mean ± SEM. *p<0.05. **p<0.01 versus SR-BI^fl/fl^ mice. #p<0.05 versus AlbCreSR-BI^fl/fl^ 0h. ##p<0.01 versus AlbCreSR-BI^fl/fl^ 0h.

### Probucolpretreatment normalizes cholesterol levels and improves AlbCreSR-BI^flf^ mice survival rate in sepsis

Serum total cholesterol (Fig4. A) and free cholesterol levels (Fig4. B) decreased after 3 days probucol diet in both AlbCreSR-BI^fl/fl^ mice and SR-BI^fl/fl^ mice. The free cholesterol level of AlbCreSR-BI^fl/fl^ mice reduced from 80.4 ± 4.0 mg/dl to 25.9 ±1.6 mg/dl after 3-days probucol feeding. The probucol diet increased the survival rate of AlbCreSR-BI^fl/fl^ mice from 14.3% to 88.9% during sepsis. But there is no difference of septic survival between SRBIfl/fl with normal diet and SRBIfl/fl fed with probucol diet (Fig4. C)

**Figure 4.**
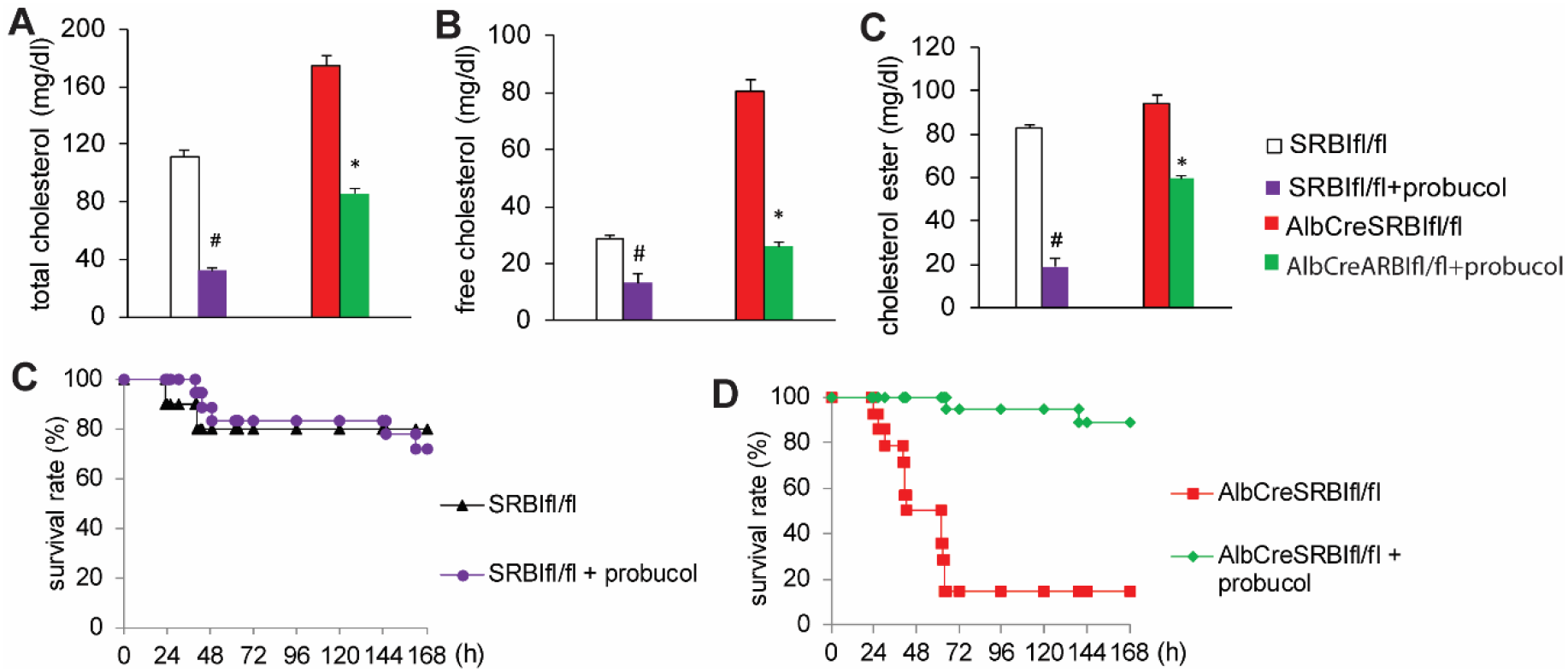
0.5 % probucol diet normalized cholesterol level and improve AlbCreSR-BI^fl/fl^ mice survival in sepsis. A, serum total cholesterol level and B, free cholesterol level of AlbCreSR-BI^fl/fl^ and SR-BI^fl/fl^ mice were analyzed at the baseline, after 3 days probucol feeding and 7 days post CLP. C, 3 days probucol pretreatment has little impact on SRBIfl/fl mice survival during sepsis but D, 3 days probucol pretreatment improves AlbCreSR-BI^fl/fl^ mice survival during sepsis. 2-tailed Student’s t-test was used to compare two groups. n=12-18 each group, mean ± SEM. *p<0.01 versus RD.

## Discussion

From our study by comparing AlbCreSR-BI^fl/fl^ mice with SRBIfl/fl mice, we come up with a conclusion that SRBI in the liver plays a markedly protective role in reducing CLP-induced septic death. SRBI expression in the liver is a key factor affecting reverse cholesterol transport and regulating cholesterol homeostasis. However, using a hypo-AlbCreSR-BIfl/fl mouse model, Huby’s group showed that the liver-specific SR-BI KO mice are not more susceptible to CLP-induced sepsis(19). An issue with their animal model is that the floxed SR-BI mice are hypomorphism (control SR-BIfl/fl mice have 90% whole body depletion of SR-BI). Huby’s group generated SR-BIfl/fl mice by flanking exon 1(23), however, the mice displaced hypomorphism due to disruption regulatory elements around exon 1. In this study, we generated SR-BIfl/fl mice by flanking exon 2. Our data shows our liver-specific SR-BI KO mice had 4-fold increase fatality than the control mice with exacerbated inflammatory cytokine production in sepsis. These results are consistent with our previous study that all confirm the conclusion of hepatic SR-BI protecting sepsis despite Huby’s group proposed the opposite argument.

Previous studies showed that disorders of lipid metabolism were critical issues in sepsis patients(24). HDL plays a protective role in sepsis and reconstituted HDL may be a therapeutic approach for sepsis(25–28). Cholesterol is the main component of these lipoproteins and be regulated during sepsis. Statin is a common drug which can lower cholesterol level by inhibiting hydroxy-3-methyl glutaryl-coenzym e A reductase (HMGCR) which is the key enzyme for cholesterol biosynthesis. In one prospective observational cohort study, patients with acute bacterial infection were divided into statin-treated group and non-statin group to observe their outcomes. Severe sepsis developed in only 2.4% of the statin group but in 19% of patients in the non-statin group(29). This indicates cholesterol is an important lipid involved in sepsis.

Sepsis and infection are associated with hemolysis and made red blood cell-free hemoglobin released into the circulation(30,31). In clinical study, sepsis caused hemolysis that massive hemoglobin release from red blood cells is a risk factor of septic death(30,32). Various potential mechanisms were considered to cause hemolysis in sepsis: 1) toxins of pathogens induce hemolysis 2) fibrin strands generated by septic coagulation can induce schistocytes 3) the complement system can impair erythrocytes viability 4) LPS and lipid binding to erythrocyte membrane lead to erythrocyte deformability(33) 5) sepsis induces eryptosis(34). Because hepatic SRBI plays a vital role in RCT to regulate cholesterol hemostasis via biliary cholesterol secretion. Deficiency of RCT causing free cholesterol accumulation in the blood leads to erythrocyte membrane cholesterol accumulation through nonspecific mechanisms and decreasing membrane deformability(35). Because of the free radicals originate from perfusion problem tissue and osmotic fragility during sepsis, the red blood is susceptible to rupture. Comparing with control mice, AlbCreSR-BIfl/fl mice showing excessive serum hemoglobin levels at CLP 20h but not at CLP 4h indicates that RCT deficiency causes aggravated hemolysis after sepsis onsite. In clinical, studies show that elevated concentrations of plasma free hemoglobin are associated with an increased risk of death and free hemoglobin is considered as a predictor of survival in severe sepsis(30,32), moreover, levels of plasma cell-free hemoglobin are independently associated with poor clinical outcomes and death, according to the use of Hemoglobin-based blood substitutes(36). Free hemoglobin can scavenge nitric oxide leading to vasoconstriction in vascular beds (37), cause lipid peroxidation and endothelial damage, regulate neutrophils (38), affect Toll-like receptor pathway and TNF-α generation(39). Also, hemoglobin is a bacterial LPS binding protein that may enhance LPS activity(40) and may involve in the production of tissue factor associated with disseminated intravascular coagulation during sepsis(41). These together let hemolysis occur on RCT deficiency mice and aggravate death during sepsis(36). After we use probucol as a cholesterol-lowering drug to decrease the abnormal cholesterol levels in RCT deficiency mice, RCT deficiency mice mitigate hemolysis and improve sepsis survival.

In conclusion, hepatic scavenger receptor BI is a key protect effector during polymicrobial bacteria-induced sepsis and can reduces septic death. Deficiency in reverse cholesterol transport leading to lipid disorder causes hemolysis and exacerbates sepsis death. Alleviation of hemolysis may be a novel mechanism of hepatic SR-BI protection against sepsis.

