## Supplemental Table 1 for "Deficiency in reverse cholesterol transport in mice augments sepsis"

Supplemental Table S1

|  | SRBIfl/fl | | AlbCreSRBIfl/fl | | P value | P value |
| --- | --- | --- | --- | --- | --- | --- |
|  | CLP 4h | CLP 20h | CLP 4h | CLP 20h | CLP 4h | CLP 20h |
| Eotaxin | 928.12±114.8 | 528.5±123.73 | 3511.51±2334.17 | 906.52±90.42 | 0.311 | 0.035 |
| G-CSF | 34021.27±2612.96 | 33546.33±6754.84 | 32228.32±1701.55 | 37360.03±1934.56 | 0.578 | 0.607 |
| GM-CSF | 48.28±22.59 | 20.04±2.8 | 20.48±2.4 | 17.54±3.9 | 0.266 | 0.615 |
| IFNy | 48.54±39.97 | 7.46±2.66 | 3.85±0.98 | 7.83±1.53 | 0.306 | 0.909 |
| IL-1a | 667.26±96.88 | 471.27±87.1 | 651.87±85.87 | 521.16±36.71 | 0.907 | 0.615 |
| IL-1B | 40.06±30.64 | 9.4±3.61 | 45.98±35.67 | 13.54±1.4 | 0.902 | 0.323 |
| IL-2 | 42.81±23.36 | 14.51±3.29 | 9.14±1.6 | 14.48±4.42 | 0.2 | 0.996 |
| IL-3 | 11.67±7.31 | 3.44±1.7 | 2.31±0.49 | 1.99±0.57 | 0.248 | 0.449 |
| IL-4 | 3.59±2.02 | 1.22±0.67 | 2.25±1.44 | 1.13±0.45 | 0.599 | 0.906 |
| IL-5 | 402.58±74.35 | 33.76±12.78 | 559.27±106.08 | 33.31±10.17 | 0.252 | 0.979 |
| IL-6 | 12116.14±3503.55 | 668.62±192.79 | 16532.15±4132.36 | 6048.92±4260.83 | 0.431 | 0.263 |
| IL-7 | 28.42±11.57 | 7.17±1.52 | 8.18±1.67 | 15.11±9.55 | 0.132 | 0.447 |
| IL-9 | 127.47±25.34 | 161.01±34.38 | 104.19±18.1 | 120.99±13.55 | 0.471 | 0.318 |
| IL-10 | 604.92±170.32 | 325.81±152.05 | 459.42±124.86 | 245.72±186.83 | 0.505 | 0.747 |
| IL-12 | 100.28±35.32 | 52.73±14.78 | 36.56±9.1 | 24.98±5.75 | 0.125 | 0.127 |
| IL-12 | 213.23±161.57 | 36.26±15.2 | 89.86±48.31 | 48.21±18.29 | 0.488 | 0.627 |
| IL-13 | 280.52±113.66 | 230.23±100.22 | 133.25±24.27 | 85.05±14.86 | 0.248 | 0.209 |
| IL-15 | 561.24±317.01 | 187.52±27.11 | 176.42±23.84 | 175.85±51.45 | 0.271 | 0.846 |
| IL-17 | 48.82±8.38 | 5.35±1.57 | 122.81±38.54 | 11.09±3.39 | 0.106 | 0.168 |
| IP-10 | 335.58±139.1 | 80.37±31.76 | 201.89±73.21 | 154.79±32.85 | 0.417 | 0.134 |
| KC | 14566.26±3022.17 | 5346.43±1691.25 | 15509.23±3086.91 | 5054.74±2263.29 | 0.831 | 0.92 |
| LIF | 8.7±1.72 | 3.05±0.42 | 11.44±2.8 | 8.99±3.87 | 0.424 | 0.187 |
| LIX | 3601.52±1268.09 | 2085.57±1389.88 | 5746.47±2062.01 | 5858.41±729.93 | 0.396 | 0.045 |
| MCP-1 | 311.39±60.84 | 79.83±27.77 | 587.28±153.63 | 143.33±44.41 | 0.134 | 0.258 |
| M-CSF | 1713.43±1414.52 | 126.95±63.6 | 49.31±13.9 | 77.34±21.85 | 0.284 | 0.488 |
| MIG | 54.06±9.56 | 31.35±13.7 | 35.22±7.34 | 35.46±12.97 | 0.146 | 0.832 |
| MIP-1a | 338.62±29.83 | 387.17±116.48 | 303.02±60.38 | 284.9±70.6 | 0.613 | 0.474 |
| MIP-1B | 767.33±149.49 | 182.56±21.49 | 569.21±118.96 | 206.9±76.32 | 0.321 | 0.77 |
| MIP-2 | 956.25±268.55 | 235.63±41.12 | 722.63±212.49 | 224.02±36.02 | 0.509 | 0.836 |
| RANTES | 171.24±81.84 | 59.46±21.46 | 41.09±5.28 | 77.41±32.4 | 0.163 | 0.656 |
| TNFa | 46.26±12.49 | 30.15±6.14 | 52.64±17.94 | 50.78±13.13 | 0.776 | 0.197 |
| VEGF | 2.67±1.38 | 0.86±0.09 | 1.7±0.67 | 1.16±0.23 | 0.541 | 0.258 |
